# A development-specific POLX essential for programmed genome rearrangement in *Paramecium tetraurelia*

**DOI:** 10.1101/2024.07.18.604053

**Authors:** Baptiste Verron, Olivier Arnaiz, Coralie Zangarelli, Nathalie Mathy, Mireille Bétermier, Julien Bischerour

## Abstract

During the sexual cycle, programmed genome rearrangement (PGR) in *Paramecium tetraurelia* involves the non-homologous end joining (NHEJ) DNA repair pathway to eliminate specific germinal Internal Eliminated Sequences (IESs) from the newly developing somatic nucleus. In addition to the core NHEJ factors Ku70/80 and Xrcc4/Lig4, additional enzymes are required to process the 4-base 5’-protruding ends generated following DNA cleavage at IES boundaries, prior to their ligation. Here, we report that PolX (a,b,c,d), four *P. tetraurelia* distant orthologs of the human Polλ DNA polymerase, are involved in repair of IES excision junctions. During rearrangements, PolX-depleted cells accumulate genome-wide errors, such as unrepaired double-strand breaks, 1-nucleotide deletions and IES retention. Although all PolX paralogs can process DNA ends, two of them (PolXa&b) are induced during PGR and have acquired tight nuclear anchoring properties through their N-terminal region, which contains a predicted BRCT domain. Finally, we show that PolXa accumulates in nuclear foci together with other NHEJ proteins and the Dicer-like enzyme Dcl5, which is involved in the biogenesis of IES-specific small RNAs. We propose that these “DNA repair foci” correspond to the sites where IES concatemers, a by-product of IES excision, are ligated together to produce the precursors of iesRNAs.

## INTRODUCTION

Programmed DNA elimination is an extreme form of genomic regulation that is well illustrated in ciliates, where two types of nuclei coexist in the same cytoplasm (nuclear dimorphism)^1^: the somatic macronucleus (MAC) is essential for gene expression but is destroyed during each sexual cycle, while the germline micronucleus (MIC) undergoes meiosis, karyogamy and copies of the zygotic nucleus differentiate to regenerate the germline and form a new somatic nucleus. Extensive genome rearrangements and multiple rounds of genome endoreplication take place during new MAC development. In *Paramecium tetraurelia,* large genomic regions and several thousands of short Internal Eliminated Sequences (IES) are eliminated via programmed genome rearrangements (PGRs)^2,3,4^. The reaction proceeds through a cut and close mechanism, in which the PiggyMac/PiggyMac-like complex (Pgm/PgmL) catalyses two precise DNA cuts centred on a TA dinucleotide that is present at the boundaries of IESs, followed by the ligation of both flanking ends by the classical non-homologous end joining pathway (cNHEJ or NHEJ)^2,5,6^. Because many essential genes are interrupted by IESs, the precision of the rearrangements is essential for the formation of a functional new MAC. It is partly achieved through the coupling between DSB introduction and DNA repair, which favours faster recruitment of repair proteins and maintenance in close proximity of the correct ends to be ligated^7,8^. In addition to the repair of the MAC junction, the NHEJ system is also used to circularise (for the longest) and assemble (for the shortest) already excised IESs into concatemers^9,10^. These concatemers provide a template for the transcription of double-strand RNA molecules and the Dcl5-dependent processing of 26-30bp short single-strand RNA (iesRNA) implicated in the epigenetic control of IES elimination^9,11^.

The *Paramecium* genome encodes NHEJ repair proteins, such as Ku70, Ku80, Ligase4 (Lig4) and Xrcc4, as well as putative homologs of DNAPKcs and Xlf ^2,12,13^, which form the minimal core machinery for the precise ligation of two compatible DNA extremities in eukaryotes^14^. In addition, the 4-base 5’ overhangs generated by Pgm cleavage are rarely compatible for direct re-joining and the final ligation step requests prior processing of the ends, including removal of one base at the 5’ end, pairing of the TA dinucleotides of each overhang and filling of the nucleotide gap^12,15^. Lessons from the study of NHEJ in other organisms has shown that processing enzymes provide NHEJ with numerous means to resolve end incompatibilities. These include NHEJ-specific DNA polymerases of the X-family (e.g. Pol λ, Pol μ), which are required to fill in overhangs and gaps, and nucleases to remove extra or damaged nucleotides (e.g. Artemis, ExoI, TdP)^14^. DSB processing enzymes are not part of the NHEJ core complex, but are recruited either of thanks to dedicated motifs like the Ku binding motif (KBM) or the BRCT domain, which mediate interaction with Ku70/80 and the Lig4/Xrcc4 complex^16,17^. The NHEJ process rarely gives rise to a unique DNA repair product when DNA extremities are not initially compatible^18^. However, the extent of end processing is minimised in favour of the ligation, by coordinating end processing with synapsis, where DNA ends are aligned^18^. In the exceptional situations of V(D)J recombination, the repair of DSBs is also intended to increase diversity at the repair site, and a specialized X-family DNA Polymerase (terminal DNA transferase or TdT), has to be recruited to add random nucleotides to the 3′-OH ends, which increase the variability of the non-templated nucleotide (N) region of the recombined gene segments^19^. By opposition, what is remarkable during IES elimination is how the 5’-end trimming and gap-filling steps are controlled to produce systematically one and only one DNA repair product. The *Paramecium* genome went through several whole genome duplications (WGDs), leading to paralogous genes from a single ancestral, which contributed to the specialisation of DNA-repair *KU80* (*KU80c*) gene for PGRs. Did specific processing enzymes emerge in a similar way during evolution, explaining the nature of end repair in PGRs?

In the present study, we have identified four *Paramecium* DNA polymerase X paralogs (*POLX a,b,c,d*) that share homology with the Pol λ identified in mammals. We show that they are involved in the gap-filling step during IES elimination and we report a singular and obligate hierarchy of end processing steps. In addition, we show that specific *POLX* paralogs (*POLXa&b*) have evolved their N-terminal domain including the BRCT, which tighten their anchoring in the new developing MAC. Finally, this work allows us to highlight a specific subnuclear structure containing the NHEJ and iesRNA synthesis factors, where we propose that IES concatemers are assembled.

## RESULTS

### The *Paramecium* genome encodes four λ-type *POLX*

A survey of genes encoding *POLX* in the *Paramecium* genome identified four paralogs that share sequence homology with mammalian *Pol λ and Pol Β (ref Antonin* ^20^). It is consistent with the fact that *λ* type of PolX is the most conserved X-family DNA *pol* throughout evolution^21^. The four genes are paralogs of an ancestral copy that was duplicated during the last two whole genome duplications (WGD) that took place during the course of evolution^22^. *POLXa* and *POLXb*, are paralogs from the last WGD, whereas *POLXc* and *POLXd* are paralogs of *POLXa* and *POLXb* from the “intermediate” WGD (Figure 1A). The four gene products have a similar size (616 aa) and are composed of a DNA polymerase domain at their carboxy terminal extremity, an 8kDa domain and a BRCT domain separated by a linker region known as S/P rich in human (Figure 1A). The presence of a BRCT domain, makes *Paramecium POLX* genes probable homologs of the *POL λ* found in mammals (Ref Papier Antonin), which together with Pol μ, is the main DNA polymerase implicated in NHEJ. PolXa share strong identity with b (91,7%), but have diverged from PolXc and d, 72,4% and 71,6 % respectively. Remarkably, the sequence of the three domains has diverged to different extent. Between the PolXa and PolXc paralogs, the 8kDa and the DNA polymerase domains are better conserved than the BRCT domain (64,5% identity) and the linker sequences (56,5% identity) (Figure 1A). *PolX* gene homologs are also present in other *Paramecium aurelia* species, where both a-type and c-type paralog can be found (Supplementary Figure S1)^23^. The time-course of gene expression during autogamy, based on RNA seq data^24^, shows that *POLXa* and *POLXb* genes are induced at the developmental stage of autogamy, whereas *POLXc* and *POLXd* are expressed constitutively at low levels during vegetative life and sexual processes (Figure 1B). The maximum of *POLXa* expression correlates with those of *PGM* and *KU70a/80c* at the DEV1 stage of autogamy (5 hours after T0), when IES excision take place in the new developing MAC ^25,26^. The presence of four *POLX*, with development-specific expressed paralogs, resembles the evolutionary scenario of the DNA repair *KU80* genes in which the *KU80c* paralog has specialized for PGRs^7^.

**Figure 1.**
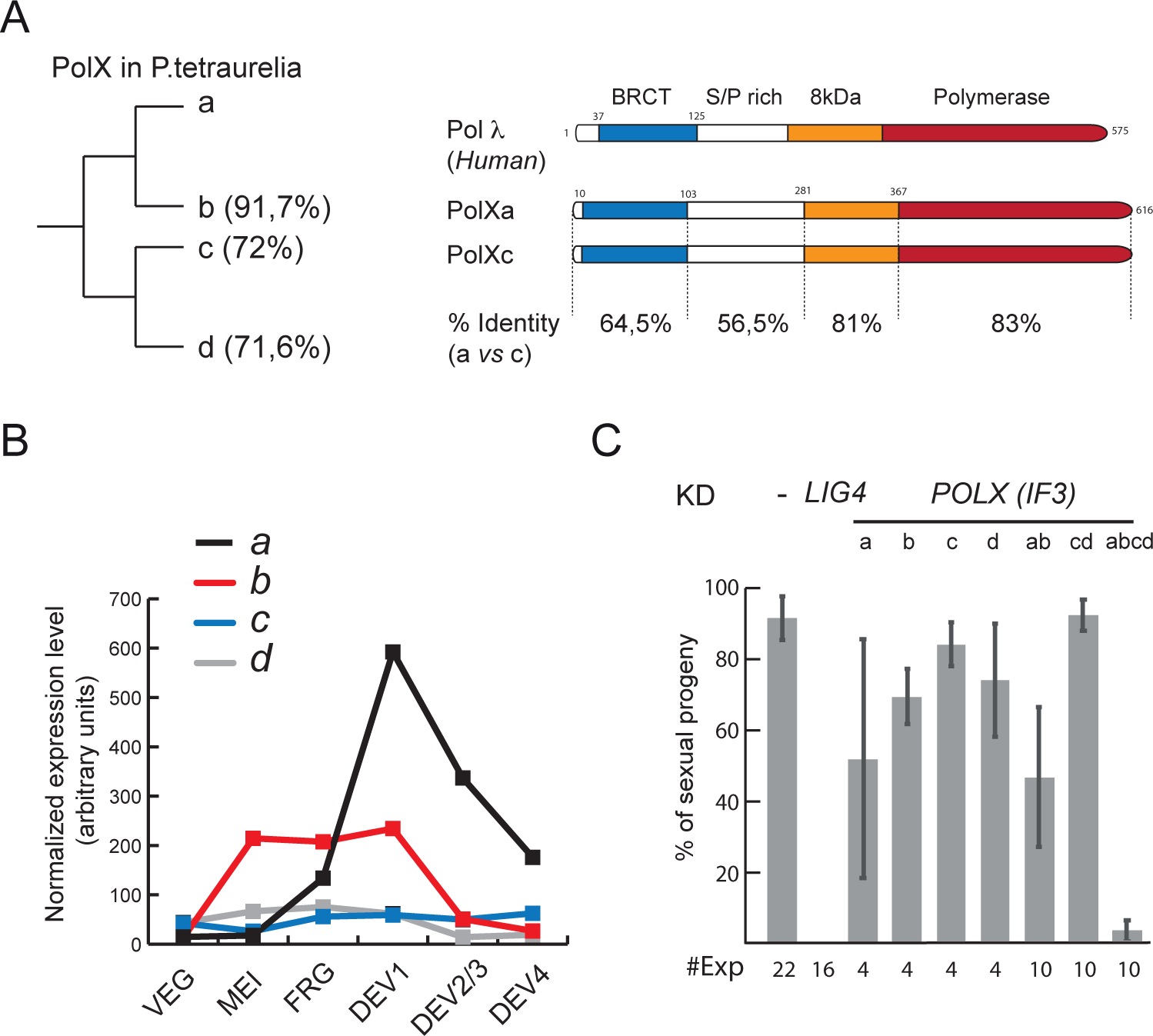
Characterisation of *P.tetraurelia POLX*. (A) Domain organisation of *P.tetraurelia* DNAPolX and diagram of the WGD relationships between *POLX* genes. The % of identity (a.a) with respect to PolXa is indicated (PolXa (PTET.51.1.P0210235), PolXb (PTET.51.1.P0360066), PolXc (PTET.51.1.P0460033), PolXd (PTET.51.1.P1010039)). The scheme of Pol λ and *P.tetraurelia* PolX domains is presented on the right. The position of the BRCT (blue), the 8kDa (orange) and polymerase (red) domains, based on sequence alignment and AlphaFold2 prediction, are depicted on the *P.teraurelia* PolXa and c homologs. The percentage of identity (a.a.) between a and c homologs is indicated for each individual domain. (B) Normalized expression level (arbitrary units) of *POLX* during vegetative cell growth and throughout the sexual process of autogamy^24^. (C) Survival test of sexual progeny in *POLX* RNAi. The percentage of viable sexual progeny is indicated with the standard deviation and the number of experiments which was performed. IF3 are feeding inserts that target their cognate gene and its closest paralog (last WGD). Co-silencing of *POLX* homologs was performed by mixing equal volumes (same OD) of feeding media. The design of the inserts and the survival tests performed with IF1 is presented in Supplementary Figure S1B.

### Co-silencing of the four *DNAPOLX* triggers death of the sexual progeny

In order to address the involvement of PolX during PGRs, we performed the knockdown (KD) of the genes, individually or in combination, by feeding *Paramecium* cells with double strand RNA producing bacteria (“feeding” technic)^27^ (Figure 1C & Supplementary Figure S1). We reasoned that the abolition of PolX activity would compromise the DNA repair step of IES elimination, which in turn would conduct to the absence of a functional new MAC and death of the sexual progeny, as we reported previously for other components of the NHEJ machinery^8,12^.

For each POLX paralog, we tested two inserts which target different sequences of the ORF (Supplementary Figure S1). The first set of inserts (IF1) were designed to silence specifically each gene, whereas the second set of plasmids (IF3) can target the cognate gene and its closest paralogs (a together with b and c together with d). The silencing of *POLX* genes individually using the specific feeding inserts (IF1) had no remarkable phenotype (Supplementary Figure S1). Similarly, mixing both IF1a and IF1b media or IF1c and IF1d media did not impact the survival of the progeny (Supplementary Figure S1). The use of the IF3 targeting *a&b* genes, either individually or in combination, leads to 30 to 50% of death in the progeny (Figure 1C). However, only the KD of the four genes, which we will refer to as *POLX* KD, performed by mixing the four IF3 a, b, c and d media lead to complete death of the progeny. We conclude from these combinatory experiments, that the developmentally induced *POLXa&b genes* are implicated in NHEJ during PGRs. However, the absence of complete death in condition of KD against *POLXa&b* suggested that *DNAPOLX* c&d genes, which are expressed constitutively at low level, are able to substitute for *POLXa&b* during PGRs.

### Transgenic *POLXc* expression restores progeny survival of PolX-depleted cells

To confirm the specificity of the quadruple KD and to definitively test the ability of both a and c-type PolX to fulfil the DNA repair function during PGR, we used a trans-complementation assay (Figure 2A). We knocked down endogenous *POLX* by RNAi, and we provided *Paramecium* cells with *FLAG-POLXa* or *FLAG-POLXc* by microinjection of their respective RNAi-resistant transgenes, in order to test their ability to complement *POLX* KD cells (Figure 2A). Both genes were placed under the control of the *POLXa* regulatory sequences, which is induced during MAC development. We tested two different tagged versions (N-ter and C-ter fusion), but only the N-terminal fused protein was functional (Supplementary Figure S2), in agreement with previous studies and structural data that demonstrate that the carboy-terminal residues of Pol λ is embedded into the catalytic site^28^. A 75kDa product, corresponding to Flag-PolXa and Flag-PolXc is detected by western-blot 5 hours (T5) after the beginning of autogamy (T0 = 50% of the cells with fragmented MAC), in agreement with the expression time course of endogenous *POLXa* (Figure 1B). Overall, both *POLXa* and *POLXc* transgenes expression correlates with their copy number determined by qPCR (cphg = copies per haploid genome). However, at the protein level, FLAG-*POLXc* is better expressed than FLAG-*POLXa* (Figure 2B & Supplementary Figure S2). Survival tests of the progeny after four days of starvation demonstrate that both PolXa and PolXc complement the depletion of the endogenous PolX. However, PolXa complements at protein level close to the limit of detection, whereas PolXc fails to complement unless higher protein level are reached (Figure 2B & Supplementary Figure S2), suggesting that PolXa is somehow more efficient than PolXc.

**Figure 2.**
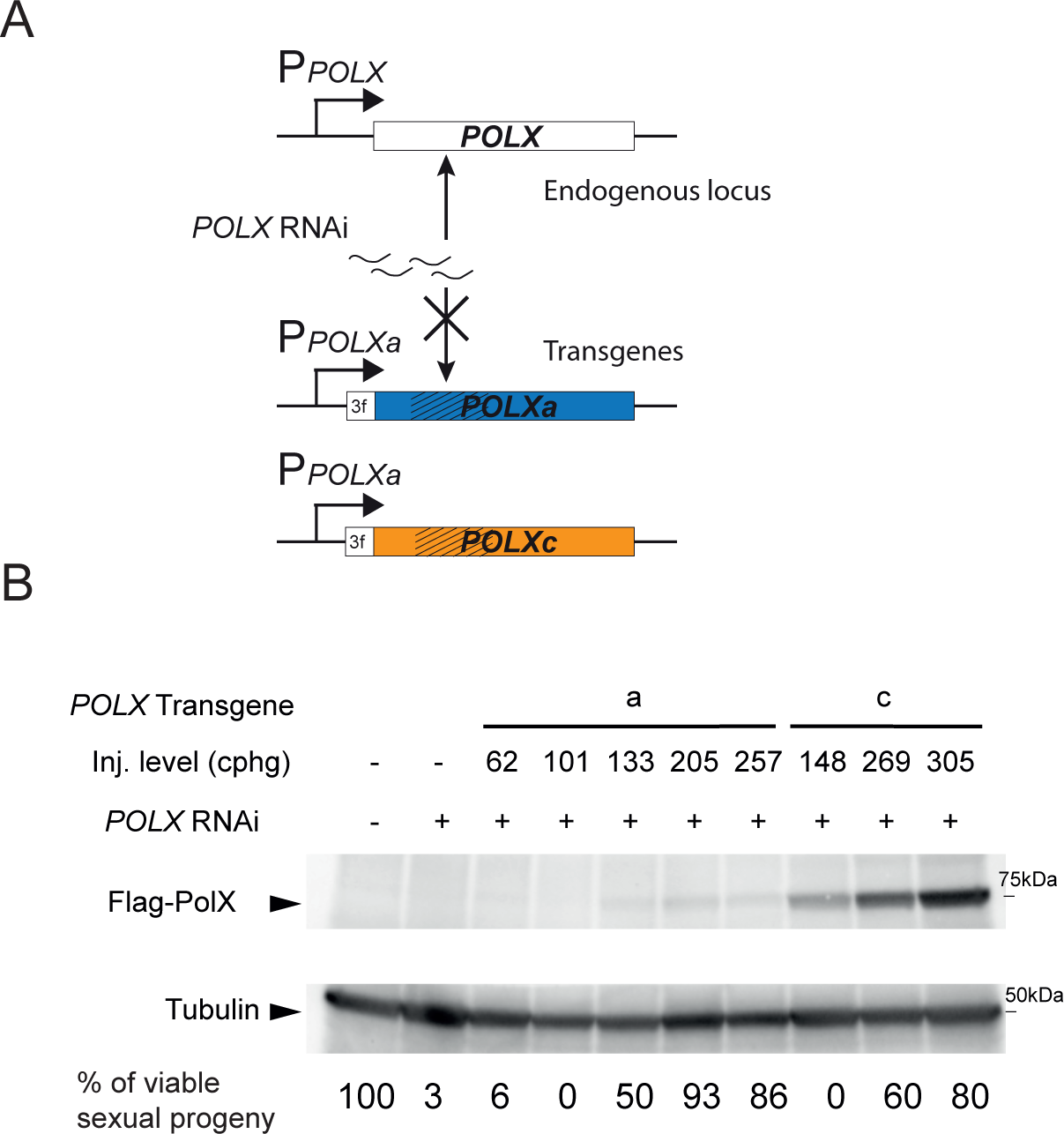
Both *POLXa* and *POLXc* complement *POLX* KD. (A) Schematic representation of transgenes used in the complementation assays. The modified *POLXa and c* nucleic acid sequences that confer resistance to RNAi are represented by hatched lines. The *FLAG-POLXa* and *FLAG-POLXc* transgenes are expressed under the control of the endogenous *POLXa* regulatory sequences. (B) Western blot analysis of *FLAG-POLXa* and *FLAG-POLXc* expression in early autogamous cells (T5) subjected to *POLX* RNAi. The injection level is indicated as copies per haploid genome (cphg). The percentage of survival of the sexual progeny is indicated for each transformant.

### PolX depletion compromises DNA repair and induces IES retention

To characterize the consequences of *POLX* KD on PGRs, cells were grown in conditions of RNAi against all *POLX*, *POLXab* or *POLXcd*. New developing MAC, collected after IESs excision has been completed under control conditions (T25), were sorted by FANS (Fluorescence assisted nuclear sorting) using antibodies directed against Pgml1^29^. The status of IES excision was then analysed using the MEND module of PartIES^8^, which rely on DNA reads overlapping IES excision sites (Figure 3A). Following *POLX* RNAi, only 46% of the reads showed correct MAC-IES junctions, while 40% corresponded to retained IESs (IES+) (figure 3A). By opposition, no IES+ reads were observed in the control or *POLXcd* KDs, as expected for the normal course of IES elimination during autogamy. To decipher which IESs are retained and excised in the absence of PolX, we calculated IES retention scores. The IES+ reads are not distributed evenly among IES, some IESs are strongly retained (IRS > 0.5) and others are completely eliminated (IRS close to 0) (Figure 3B and Supplementary Figure S3). Groups of IESs were previously defined according to their excision timing during development^29^. This excision timing correlates with the distribution of retention scores in *POLX* KD, with very early excised IES being almost all excised (0 < IRS < 0,2), whereas late IES are almost all retained (0,7 < IRS < 1) (Figure 3B). We observed the same trend for a *LIG4* KD, where DNA-repair defaults happening during excision of the first IESs inhibit excision of latest IESs^8^.

**Figure 3.**
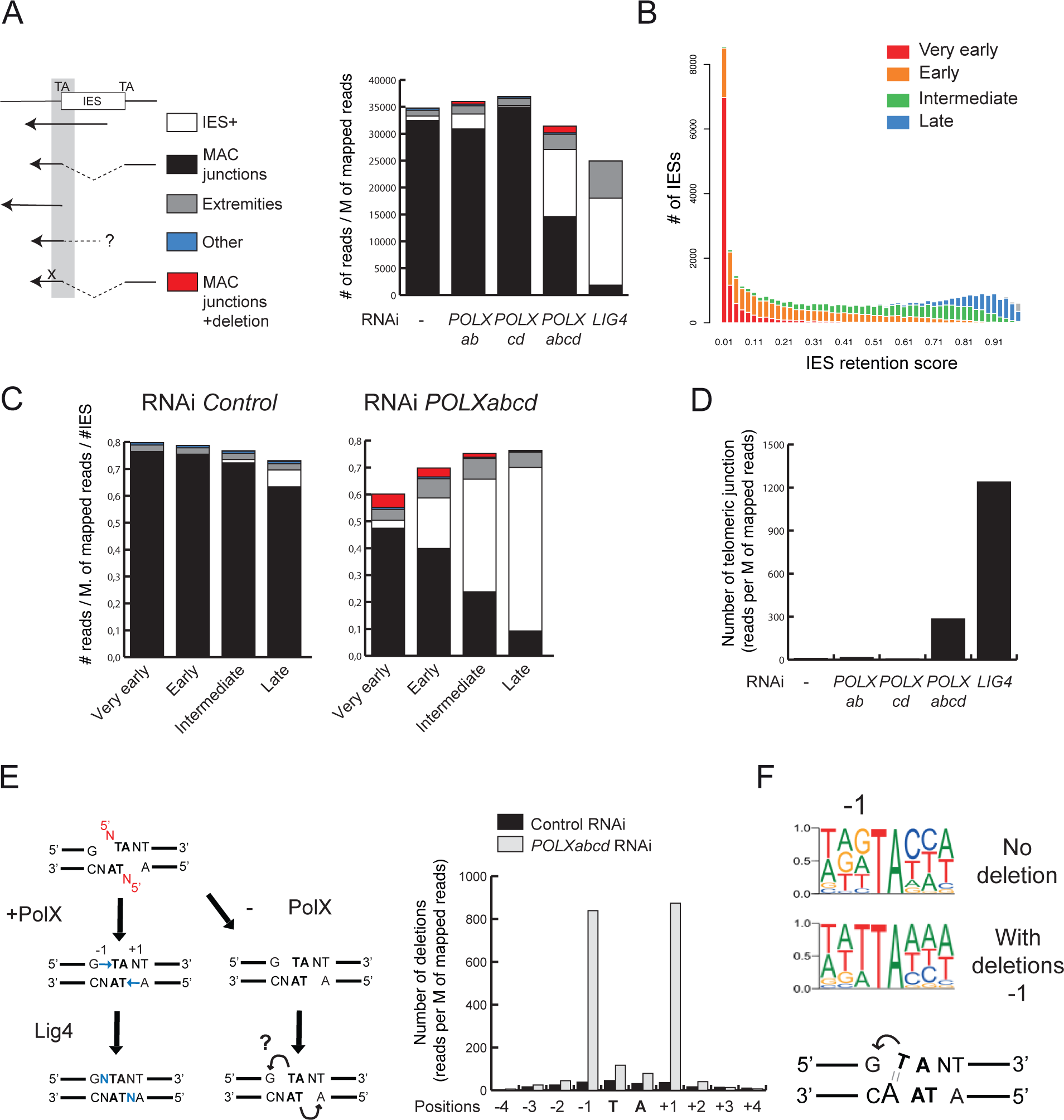
Genome wide analysis of rearrangements in PolX depleted cells. (A) Analysis of the reads covering IES excision sites in control and *POLX* KD conditions^8,39^.The *Illumina* reads, represented by black arrows, are classified in five categories depicted on the left. They correspond to (i) the genomic loci with an IES (IES+), (ii) excised IES (MAC junction), (iii) reads starting at the TA (+ or −2 bp) (“Extremities”), (iv) reads mapping partially on the flanking DNA but stopping on the TA dinucleotide (+ or −2 bp), (v) MAC junction reads with a deletion flanking the TA dinucleotide (MAC junction + deletion). Lig4-depleted cells data are extracted from (*Bischerour et al,* 2023)^8^. (B) Histogram representation of IES retention score (IRS) distribution. The excision timing profile to which IESs belong is indicated. The IESs in grey are undetermined for their excision timing profile. (C) MEND analysis of IES excision timing clusters in control and *POLX* KD cells. The number of reads is normalised by the number of IESs in each cluster. (D) Abundance of *de novo* telomere addition sites in different *POLX* KD conditions. As previously described ^29^, a telomere addition site was pinpointed whenever a read alignment stops on the MAC sequence and the read sequence proceeds with telomeric repeats (G_4_T_2_ or G_3_T_3_). The number of such telomere addition events was normalized by the total number of mapped reads (cpm). (E) Scheme of ends processing scenario in presence or absence of PolX. The arrows represent a ligation trough the gap which will generate of deletion at the −1 an +1 positions. (F) Frequency of reads containing a deletion around TA excision sites in control and *POLX* KD. (G) Logos of base frequency for IES excision site either with (>5 reads) or without deletion (0 reads) at the −1 positions. Hypothetic scenario of ligation through the gap when an adenine is located opposite to the gap. IESs retention score, MEND analysis of IES excision time clusters and quantification of deletions at −1 and +1 positions for *POLXab* and *POLXcd* KD are presented in Supplementary Figure S3.

In addition to IES retention, we observed like in *LIG4* KD, a decrease of read coverage around IES excision sites (Figure 3A). We also noted a slight increase of reads “extremities” which correspond to reads finishing at the TA position of IES excision sites (Figure 3A & S3). These two phenotypes are particularly conspicuous for the IESs of the two earliest time group of excision time (Figure 3C and S3), which are excised, supporting a scenario of unrepaired broken ends when PolX is depleted. Concomitantly, *de novo* telomere addition, which constitute an alternative DNA repair outcome when end-joining repair is not effective is detected in PolX depleted cells (Figure 3E). Taken together, we conclude that depleting the four PolX paralogs leads to the accumulation of unrepaired DSBs at the excision site of the earliest IESs, which impairs the elimination of the following IESs.

### PolX depleted cells accumulate DNA repair errors at IES MAC junctions

In condition of *POLX* KD, many sequencing reads mapping to new MAC junctions (IES-) contain a deletion of one nucleotide close to the TA of IES excision sites (in red) (Figure 3A, 3C, 3E and Supplementary Figure S3). By collapsing the reads showing a deletion, we mapped almost all deletions to the −1 and +1 positions, corresponding to the positions that are filled in by PolX during DSB repair (Figure 3E). This phenotype is specific of the *POLX* KD because it was not observed in other KDs conditions^8^. One hypothesis to explain the presence of a gap, is a ligation reaction through the nucleotide gap, which is likely to persist in PolX-depleted cells (Figure 3E). Interestingly, at the level of an IES, the deletions are often introduced preferentially if not exclusively on one side of the junction (Supplementary Figure S3). By analysing base frequencies, we tested whether the sequence surrounding the TA (+/-3 bases) differed between junctions with deletion at position −1 (> 5 reads) and without deletions (0 reads) (Figure 3F and Supplementary Figure S3). The sequence of MAC junctions having deletions are enriched for a thymine at the position of the gap, and by consequence an adenine on the opposite strand. If the deletion was caused by ligation through the gap, pairing between this adenine and thymine of the TA dinucleotide located on the opposite strand will bring the bases on either side of the gap closer together, favouring ligation (Figure 3F). The same analysis on IES excision sites with deletions at the +1 position, confirm the enrichment for an adenine opposite to the gap (Supplementary Figure S3). All together, these observations suggest that the sequences surrounding the TA is a strong determinant of deletion, and support the scenario of a ligation through the gap when PolX is missing.

Unrepaired DSBs, deletions at −1 or +1 positions and *de novo* telomere additions are DNA-repair defaults associated with the depletion of PolX. We observed the same default at lower frequency in *POLXa&b* but not in *POLXc&d* KDs (Figure 3A, 3D and Supplementary Figure S3). It provides an explanation to the lethality of the sexual progeny (Figure 1C) in these conditions of RNAi and further support a predominant role of PolXa&b during PGRs.

### IESs flanked by compatible DNA ends are equally sensitive to the depletion of PolX

A subset of IESs (6.8% of all IESs) have 4-base direct repeats at their two flanking sequences, which may permit direct ligation of the two cohesive flanking ends without the need for any end processing. We hypothesised that this group of IESs may be insensitive or less sensitive to the depletion of PolX. We focused our analysis on the subset of very early excised IESs and compared those with compatible versus incompatible DNA flanking sequences (Supplementary Figure S3). The fraction of correct MAC junction and errors doesn’t change significantly between IESs with and without compatible ends in condition of *POLX* KD. The same DNA repair defaults are detected (unrepaired DSBs and deletions at −1 and +1 positions), suggesting that PolX is also required for the repair of IES excision site with compatible flanking DNA ends.

### The N-terminal BRCT containing domain strengthen PolXa anchoring in the new developing MAC

To further characterise the a- vs c-type PolX enzymes, we investigated their localisation by immunolabeling of Flag-tagged proteins during autogamy. Two transformants, with similar protein expression levels of Flag-PolXa and Flag-PolXc, were analysed at T5 of autogamy (Figure 4A and Supplementary Figure S2B). The immunolabeling conditions, which include a Triton permeabilization step prior to fixation, favours the observation of stable nuclear complexes^7,30^. Whereas Flag-PolXc signal is barely detectable, Flag-PolXa signals can be detected in the newly developing MAC, where it concentrates in nuclear foci, suggesting a difference of localisation or anchoring between c and a-type PolX (Figure 4A).

**Figure 4.**
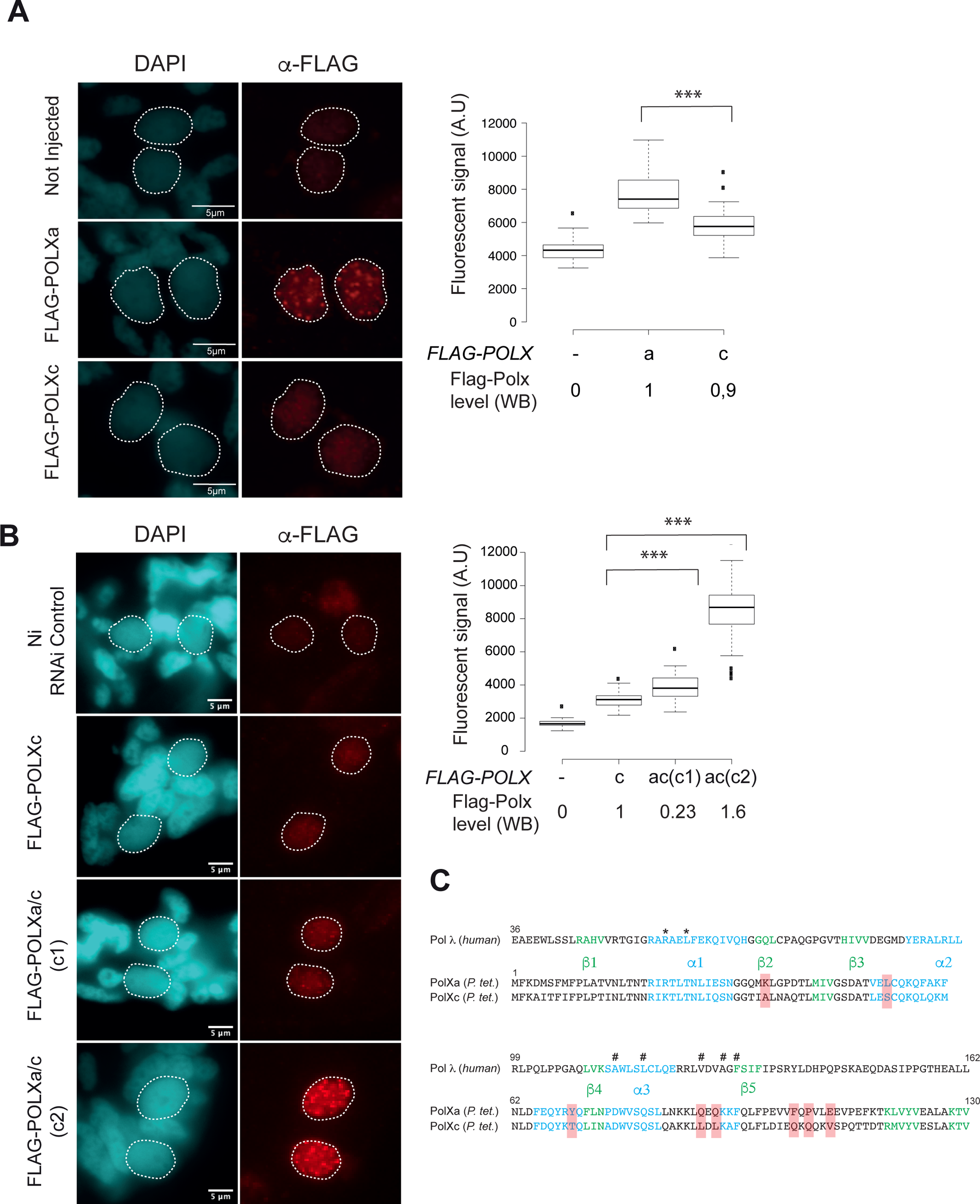
PolXa has specific nuclear anchoring properties provided by its N-terminal domain. (A) Localisation of PolXa and PolXc in the new developing MAC. The nuclear localisation of Flag-PolXa and Flag-PolXc is addressed by immunolabelling at early time of autogamy (T5-T10) in condition of *POLX* RNAi, using anti-Flag antibodies and a Triton pre-extraction step protocol^7^. The Flag mean fluorescence intensity in developing MACs, and the relative amount of Flag-PolXa and Flag-PolXc proteins is presented on the right. (B) Localisation of the chimeric PolXa/c protein in the new developing MAC. The Flag mean fluorescence intensity in developing MACs and the relative amount of expressed Flag-PolXa and Flag-polXc proteins quantified by western-blot are presented on the right side. The western-blot used for quantification and the detailed analysis of Flag signals and anlagen size of the experiments are presented in Supplementary Figure S4. For A and B panels, fluorescence intensity analysis, boxplot representation and statistical analysis were performed as described^34^. All numerical values that were used for the boxplot representations can be found in Supplementary Table S2. In all statistical analyses ** stands for p<0.01 and *** for p<0.0001 (Mann-Whitney-Wilcoxon statistical test). (C) Structural alignment of Polλ (residues 36-162) and *Paramecium* N-terminal domain (residues 1-130) of PolXa and c. Superposition of protein sequences, secondary structure elements determined by NMR for human Polλ^31^, and AlphaFold2.3 prediction for *Paramecium* PolXa and c. β-strands and α-helices are coloured in green and blue respectively. Conserved residues among *aurelia* species, with remarkable biochemical differences between a and c-type of PolX are highlighted in salmon. Human Polλ residues implicated in Ku/Lig4/Xrcc4 interaction are marked by asterisks. Surface-exposed hydrophobic residues identified in human Polλ and previously implicated in BRCT domain stability, are marked by a hashtag^31^.

The linker and the BRCT domain of PolX paralogs, diverged more extensively than the rest of the protein (Figure 1A). Besides, the BRCT is essential for the human Pol λ recruitment to the DNA repair complex through interactions with Ku70/80 and Lig4/Xrcc4 complexes^17^. Therefore, we speculated that a and c-type BRCT sequence divergency is at the origin of their nuclear anchoring properties. To test this hypothesis, the residues 1 to 170 of PolXc, which includes the BRCT domain (positions10-103), was replaced by that of PolXa, and the chimeric protein PolXa/c was tested for its ability to complement *POLX* KD and to anchored in the new MAC (Figure 4B & S4B). One *FLAG-POLXc* and two *FLAG-POLXa/c* transformants, respectively expressing low (0.23x) and high (1.6x) amount than PolXc were analysed at T5. The three transformants expressed sufficient PolX to produce viable sexual progeny (Supplementary Figure S4C), but signal of Flag-PolXc remained low compared to Flag-PolXa/c (Figure 4B). Moreover, the PolXa/c chimera forms foci similar to those observed for PolXa protein, suggesting that a-type N-terminal domain, is sufficient to provide PolXa with tight nuclear anchoring properties. The structure of the Pol λ BRCT has been solved and mutation located in the first α helix (α1) were shown to impaired Pol λ interaction with other NHEJ partners (residues marked by *)^17,31^. We combined Alfafold2.3 predicted structures and the primary sequence alignment of the a- and c-type BRCT domain of *P.aurelia* species to identified potential differences in protein sequences that could account for this phenotype. The secondary structure components were mapped, but no striking differences between a and c (highlighted in salmon) are observed in or in the vicinity of α1 (Figure 4C). Interestingly, a set of hydrophobic residues (marked by #) including L88 in PolXc are substituted for polar amino-acids like glutamine in PolXa. The L88 position corresponds to the evolutionary conserved V125 residue in Polλ, Polμ and TdT^31^. It locates into a hydrophobic patches of surface exposed residues which was show to be essential for BRCT domain stability^31^. Although it suggests that both a and c-type BRCT may have similar interaction surface for NHEJ partners, the way this domain is interacting or exposed to the rest of the protein might be different in the two proteins.

### PolXa foci co-localizes with Ku and Lig4

Pol λ is recruited by core NHEJ factors through its BRCT domain and previous localisation studies of *Paramecium* Ku70a and Ku80c showed a similar organisation in nuclear foci^7,26^. We tested whether PolXa and Ku foci may correspond to the same structure, using the Flag-PolXa fusion and a polyclonal antibody raised against the C-terminal peptide sequence of Ku70a to detect the endogenous protein^8^. Of note, a control immunolabelling of Ku70a-depleted cells still shows a signal above background in the developing MAC, suggesting that the antibody is not fully specific for Ku70a^32^. Nevertheless, this antibody revealed the *Flag* foci in *FLAG-KU70a* transformed cells (Figure 5A). Using *FLAG-POLXa* injected cells in conditions of *POLX* KD, we observed that PolXa and Ku70a foci colocalise (Figure 5A). Similarly, we tested the presence of Lig4, which is known to interact with Pol λ. We used cells transformed with a *FLAG-LIG4* transgene under the control of its own regulatory sequences that we grow up in RNAi control condition. We observed that the protein localises in the new developing MAC, as previously observed for a GFP-fusion^12^ but also accumulates together with Ku in the nuclear foci, which was not observed previously with GFP-fused protein, probably because of experimental setup discrepancies.

**Figure 5.**
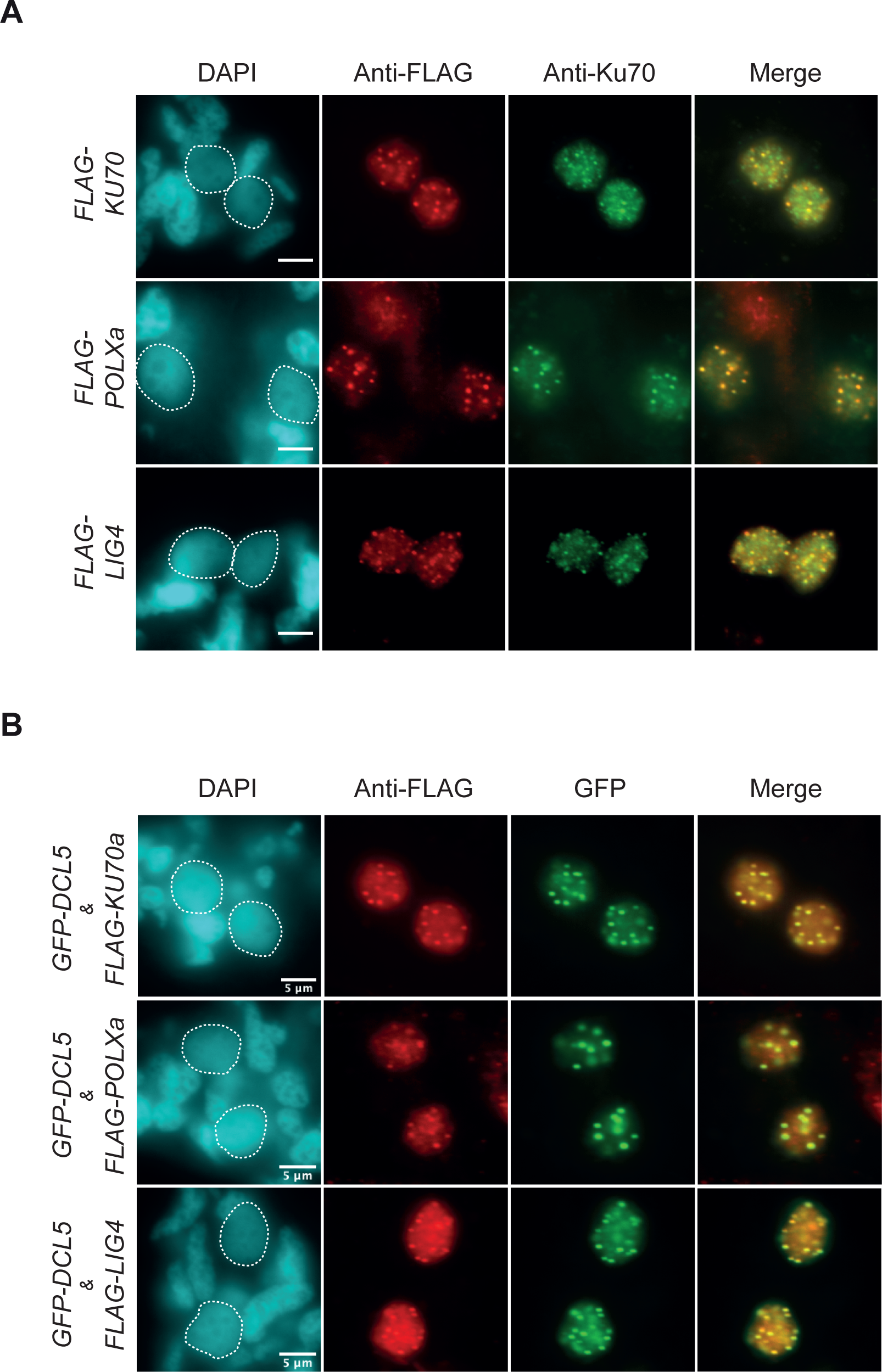
DNA repair Ku70, Lig4 and PolXa factors colocalise in nuclear foci together with Dcl5. Localisation of Flag-Ku70a, Flag-PolXa, Flag-Lig4 and GFP-Dcl5 during autogamy. (A) The nuclear localisation of Flag tagged proteins was addressed by immunolabelling of transformed cells at early time of autogamy (T5-T10). *FLAG-KU70a* and *FLAG-LIG4* injected cells were grown in control RNAi medium, whereas *FLAG-POLXa* injected cells were submitted to *POLX* RNAi. Scale bar is 5 μm. (B) Colocalisation of *GFP-Dcl5* and *Flag-Ku70a*, *Flag-DNAPolXa* or *Flag-LIG4* in early autogamous cells(T5-T10). *FLAG-KU70a* and *FLAG-DNAPOLXa* transformants were grown in condition of *KU70a* and *DNAPOLX* RNAi respectively, whereas *FLAG-LIG4* transformant was submitted to control RNAi. For all cells where foci can be observed, we confirmed the colocalisation of Flag and GFP foci.

We conclude that several DNA repair factors including Ku70/80, Lig4 and PolXa colocalise or cluster together in foci at early time of MAC development when IES excision take place.

### DNA repair factors colocalise with Dcl5 in nuclear foci

In addition to the DNA repair step that takes place at each individual IES excision site along the newly developing MAC genome, excised IES sequences are assembled into extrachromosomal concatemers, and/or circularized for the longest, through a Lig4-dependent ligation of their ends^9,12^. Following DNA repair, these assembled IESs are the substrate of iesRNA synthesis. Dcl5 is the specific Dicer-like protein involved in their biogenesis and was found to form foci in the newly developing MAC at the time of IES excision^11^. We co-transformed cells with *GFP-DCL5* and *FLAG-KU70a* transgenes and subjected the transformants to *KU70a* RNAi to knock-down the endogenous *KU70a* gene and increase the Flag-Ku70a signal^8^. We found that GFP-Dcl5 foci overlapped with Flag-Ku70a foci, suggesting that the DNA repair machinery that concentrates in these structures, correspond to regions where IES concatemers are assembled in the new developing MAC. We confirmed the result using *GFP-DCL5*-injected cells and immunolabelling of endogenous Ku70a (Supplementary Figure S5). Furthermore, we used GFP-DCL5 and *FLAG-LIG4* or *FLAG-POLXa* co-injected cells, to demonstrate that the three DNA repair factors colocalise in the same structures, together with Dcl5 (Figure 5B).

## DISCUSSION

### The localisation of PolX pinpoints two different DNA repair compartments during IESs excision

So far, our attention has been focused on the repair of chromosomal IES excision sites. We proposed previously that DSB introduction and DNA repair are tightly coupled during PGRs in *Paramecium*^7,26^. The nuclear localisation pattern of PolX and Lig4 reported here, with an evenly distributed signal in the new developing MAC together with nuclear foci, resembles the pattern observed previously with Ku70/80 proteins at early autogamy stages^7,26^. We provide evidence that PolX, Ku and Lig4 foci colocalise, suggesting that they are part of a same DNA repair “structure”. Could these foci be the nuclear compartments, in which IES boundaries are cleaved by Pgm and chromosomal MAC junctions are repaired at IES excision sites?

We observed that endogenous Pgm and a functional GFP-Pgm fusion can also form foci^7,33^. However, the colocalisation of Ku and Pgm foci is unclear and might be only partial. Moreover, Flag-tagged PgmLs, which were proposed to assemble into a higher-order excision complex together with Pgm, are essential for IES elimination but were not shown to form foci by the time IES excision takes place^34^. How to reconcile these discrepancies and propose a function for the foci containing DNA repair factors? The presence of Dcl5, the Dicer-like enzyme responsible for the biogenesis of iesRNAs^11^, in the same foci as Ku, Lig4 and PolX, provides an answer to this paradox. Indeed, IES concatemers are generated by NHEJ-mediated ligation of excised IES, and their bidirectional transcription produces double-stranded RNA molecules that constitute substrates for Dcl5-dependent cleavage. For this reason, we propose that Ku, Lig4, PolX and Dcl5 foci are the place where IESs are ligated to each other and transcribed to give rise to dsRNA precursors that are subsequently processed into iesRNAs by Dcl5. This provides an explanation for the partial or complete absence of the proteins that only contribute to DNA cleavage. The multiple rounds of DSB repair of spatially close extremities, which are required to assemble IES concatemers, may contribute to the concentration of NHEJ factors in nuclear foci. In contrast, the uniform background immunolabeling signal associated with DNA repair proteins in the new MAC may correspond to the repair of IES excision sites along the genome. Thanks to the formation of DNA repair foci, free IESs are assembled into concatemers (or intramolecular circles) at a limited number of subnuclear places, which might be on purpose. We propose that beyond its role in iesRNA biogenesis, the function of IES clustering and assembly into concatemers is to sequester IES ends and impede their potentially detrimental re-insertion into the MAC genome, which has been observed when we perturbed the system^8^.

So far, our attention was focused on the repair of the chromosomal IESs excision site. The localisation pattern of PolX and Lig4, with an evenly distributed signal in the new developing MAC together with nuclear foci resembled those observed previously with Ku70/80 protein at the early time of autogamy^7,26^. We demonstrated that PolX, Ku and the Lig4 foci colocalise, suggesting they are part of a same “DNA repair structure”. We proposed that DSBs introduction and DNA repair are tightly coupled during PGRs^7,26^. The endogenous Pgm and GFP-Pgm fusion can also forms foci^7,33^ and it is tenting to proposed that it corresponds to ongoing repair of the IES MAC junctions. However, the colocalization of Ku and Pgm foci is unclear and might be only partial. Moreover, Flag-tagged Pgmls, which were proposed to assembled into a higher order excision complex together with Pgm^34^, are essential for IES elimination, but were not shown to form foci. How to reconciled these two propositions and to explain foci containing DNA repair factor? The presence of Dcl5, the dicer-like enzyme responsible for the biogenesis of iesRNA, in the same foci as Ku, Lig4 and PolX, provided an answer to this paradox. IES concatemers are products of the ligation of excised IESs via the NHEJ machinery, and the substrate of Dcl5, which cleave double strand RNA molecules resulting from its transcription. For this reason, we propose that Ku, Lig4, PolX and Dcl5 foci are the place where IESs are ligated to each other. This provides an explanation for the partial or complete absence of the protein that only contribute to the DNA cleavage. The multiple rounds of DSBs repair of close extremities, which are required to assembled IESs concatemer will contribute to the concentration of NHEJ factors. While the even immunolabeling signal associated with DNA repair protein in the new MAC correspond to the repair of IESs excision site along the genome, the free IESs are assembled at a limited number of subnuclear places, which might be on purpose. We propose that the beyond its role in iesRNA biogenesis, the function of IES clustering and assembly into concatemer is to sequester them to impede their re-ligation into the MAC genome, which will be detrimental, but can happens when coupling between DNA cleavage and repair is perturbed^8^.

### NHEJ follows an obligate processing of IES ends during PGR

The DNA repair step involves a limited processing of the DNA ends, with the removal of one nucleotide at each 5’ end and subsequent addition of one nucleotide to fill the gap. In PolX-depleted cells, DNA repair is impaired at the ligation step, leading to the accumulation of unrepaired DSBs. However, in favourable sequence contexts of (presence of an adenine opposite to the gap), a ligation through the 1-nucleotide gap may happen generating a deletion at −1 or +1 positions relative to the TA at the MAC chromosomal junction. These deletions were not observed in other DNA-repair deficient contexts and constitute a signature of the absence of gap-filling. Unexpectedly, these errors are also observed at the excision junctions of IESs with 4-base direct repeats at their two flanking sequences, indicating that base removal at the 5’ end is systematic and probably precedes the pairing of the two 5’ protruding extremities. This is an interesting distinction from the hierarchy and kinetics of end processing in the classical NHEJ DSB repair pathway, where it has been proposed that base pairing and the ligation take precedence over end maturation^18^. When two partially compatible ends are provided for DNA repair, heterogenous DNA repair are often generated due to alternative pairing of the cohesive ends. The removal of the 5’ nucleotide abolishes possible alternative pairing of the first base and makes the two TA dinucleotides the unique solution for pairing. How the 5’ processing is restricted to the removal of only one base without the possibility to probe the compatibility of DNA ends remains an intriguing aspect of the *P.tetraurelia* NHEJ system during PGR.

### Specific Pol X paralogs contribute to NHEJ during PGR

We have identified four PolX orthologs in the *P.tetraurelia* genome that have the domain organization of the mammalian Polλ, including a BRCT domain. The ability of both PolXa and PolXc to carry out DNA repair during IES excision seemed to initially rule out a scenario of specialisation during evolution. This makes a clear difference with the Ku80 a and c paralogs, Ku80a being unable to activate Pgm-dependent DNA cleavage^35^. However, we noticed that higher amounts of PolXc than PolXa are required to get complete survival of the progeny. We also observed a marked difference in the way the two proteins are anchored in the new developing MAC. Their N-terminal domain, which includes the BRCT domain, has diverged during evolution and contributes to the specific properties of PolXa. We proposed that the BRCT domain of PolXa tightens its interaction with NHEJ factors, allowing it to perform its function at lower concentration. How the BRCT domain of Polλ and other PolX implicated in NHEJ interacts precisely with Ku and Lig4/Xrcc4 is unknown. It has been reported that mutations located in the α1 helix of the human Polλ and μ BRCT domain abolish its interaction with the NHEJ machinery^17,31^, but no clear differences were observed in this helix between *Paramecium* a and c-type PolX. Interestingly, however, a clear difference lies in a region conserved in Polλ and μ (V125 in Polλ), which is implicated together with the BRCT α3 helix, in the formation of a hydrophobic patches exposed to the surface^31^. The c-type BRCT domain have conserved hydrophobic residues at the positions L88 and L90, whereas amino acids were replaced by polar residues in PolXa (glutamine). These substitutions may change the way the BRCT domain and the linker domain of PolXa interacts and are positioned relative to other NHEJ partners. How may tightening the interaction with NHEJ factors be beneficial? In human, Polλ activity is regulated during the cell cycle, to restrict its activity during G2 and late S-phase^36^. Moreover, a mutator phenotype associated with PolX over-expression has been reported previously^37,38^. In *Paramecium*, PGRs and genome endoreplication occur simultaneously during sexual processes. For this reason, a high level of PolX expression may be detrimental, and may have driven the selection of a PolX protein with better affinity for the NHEJ factors, in order to limit interferences with the replication machineries. On the other side, the c-type PolX proteins, which are expressed constitutively, are likely to perform additional DNA repair functions, such as base excision repair (BER), for which a BRCT domain is dispensable. From this point of view, the divergence between the PGR-associated a-type PolXs and the c-type PolXs is reminiscent of the evolutionary split that gave rise to Polλ and Polβ in metazoans.

## ACKNOWLEDGMENTS

This work was supported by the *Centre National de la Recherche Scientifique* (CNRS) [intramural funding to M.B.], the French National Agency for Research (ANR) [grants number ANR-21-CE12-0019 “CURE” to M.B], the *Fondation pour la Recherche Médicale* [grant FRM EQU202103012766 to M.B.], the ARC Foundation for Cancer Research [grant number PJA_A2020_CA2020121to J.B.]. We acknowledge the sequencing and bioinformatics expertise of the I2BC High-throughput sequencing facility, supported by France Génomique (funded by the French National Program “Investissement d’Avenir” ANR-10-INBS-09). The present work has benefited from Imagerie-Gif core facility supported by I’Agence Nationale de la Recherche (ANR-10-INBS-04/FranceBioImaging;ANR-11-IDEX-0003-02/ Saclay Plant Sciences) This work was supported by the French Infrastructure for Integrated Structural Biology (FRISBI) ANR-10-INBS-0005. The authors thank Linda Sperling, Joël Acker and Vinciane Regnier for their critical reading of the manuscript. We would like to thank Antonin Nourisson, Marc Delarue and the team members for their discussions and for sharing results and data on the structure of PolX proteins.

## AUTHOR CONTRIBUTIONS

Conceptualization, Methodology,; Investigation,; Software,; Data curation, Writing – Original Draft,; Writing – Review & Editing,; Funding Acquisition,; Resources,; Supervision,

## DECLARATION OF INTERESTS

“The authors declare no competiting interests”

## MATERIALS AND METHODS

### *Paramecium* strains, culture conditions, and gene knockdowns during autogamy

*P. tetraurelia* wild-type 51 new or its mutant derivative 51 *nd7-1* were grown in a standard medium made of a wheat grass infusion inoculated with *Klebsiella pneumoniae* and supplemented with 0.8 µg/mL β-sitosterol and 100 µg/mL ampicillin. Autogamy was carried out as described ^41^ and the progression of old MAC fragmentation and new MAC development was monitored using a Zeiss Lumar.V12 fluorescence stereo-microscope, following quick fixation and staining of cells in 0.2% paraformaldehyde 20 µg/mL DAPI. The T0 time-point for each experiment is defined as the stage where 50% of cells in the population harbour a fragmented old MAC.

RNAi was achieved using the feeding procedure, as described ^34,42^. *Paramecium* cells grown for 10 to 15 vegetative fissions in plasmid-free *Escherichia coli* HT115 bacteria were transferred to medium containing non-induced HT115 harbouring each RNAi plasmid and grown for ∼4 additional divisions. Cells were then diluted into plasmid-containing HT115 induced for dsRNA production and allowed to grow for ∼8 additional vegetative divisions before the start of autogamy. Final volumes were 50 to 100 mL for middle-scale experiments (western blotting, immunostaining and DNA extraction) and 0.5 to 1 L for large-scale experiments (whole-genome sequencing). The presence of a functional new MAC in the progeny was tested after four days of starvation as described in ^12^.

Control experiments were performed using the L4440 vector. RNAi plasmids were L4440 derivatives carrying the following inserts: KU70a-1 (bp 514–813 from *KU70a*) ^43^ and pLIG4b-L(bp 521-1774 from LIG4b) for LIG4 ^12^.

### Transgene construction, micro-injection and protein expression analysis

Silent mutations were introduced into the *POLXa and c* nucleic acid sequence to make it insensitive to RNAi and 3XFLAG tag was added at the 5’ or 3’ end. Plasmid sequences of plasmids are presented in Supplementary File S1. All transgene-bearing pUC18 derivatives were linearized with appropriate restriction enzymes (Bsa*I*) and co-injected with an *ND7*-complementing plasmid into the MAC of vegetative 51 *nd7-1*. Sequences of the *FLAG-tagged* transgenes encoding N-terminal fusions of the 3X Flag tag (YKDHDGDYKDHDIDYKDDDDKT) to POLX and LIG4 are displayed in File S1. Transgene injection level (copy per haploid genome or cphg) was determined by qPCR on genomic DNA extracted from vegetative transformants, using a LightCycler 480 and the Luna qPCR Mix (New England Biolabs). Oligonucleotide primers and the genomic reference locus are listed in Table S2. GFP-DCL5 construct was kindly provided by the laboratory of M.Nowacki ^11^. All transgenes are under the control of their endogenous promotor, except for the POLXc and POLXac genes that were expressed under the control of the POLXa promotor sequences. Co-injection (GFP-DCL5 and FLAG-LIG4, KU70 or POLXa) were made with 1/3 DCL5, 2/3FLAG ratio of linearized plasmid.

A peptide corresponding to amino acid sequence 588 to 602 of *P. tetraurelia* Ku70a was used for rabbit immunization to yield a-Ku70a (588-602) antibodies (Eurogentec). Polyclonal antibodies were purified by antigen affinity purification. The FLAG tag was revealed using monoclonal anti-Flag antibody α-FLAG M2 (Sigma-Aldrich).

Protein extracts used for western-blot were prepared as previously described in ^7^.

### Cell staining

Cell staining and Immunostaining of fixed cells using polyclonal anti-Ku70a rabbit antibody α-KU70 C-ter^8,32^ or monoclonal anti-Flag antibody α-FLAG M2 (Sigma-Aldrich) were performed as described previously ^7^. Observations were made with an Olympus BX63 epifluorescence microscope with a 63x oil objective or an Olympus BX63 epifluorescence microscope with a 60x oil objective, focusing on the maximal area section of new developing MACs. Quantification of new MAC sizes, fluorescence intensity, boxplot representation and statistical analysis were performed as describe ^7^.

### Purification of new developing MACs, sequencing of genomic DNA and mapping on the *Paramecium* genomes

New developing Macs (anlagen) were purified by FANS using anti-Pgml1 antibodies and genomic DNA was extracted as previously described in ^29^. The total genomic DNA of late autogamous cells (4 days of starvation) was extracted from large-scale cultures (1litre) using the NucleoSpin Tissue extraction kit (Macherey Nagel) as previously described ^7^. All genomic DNAs were sequenced at a 76 to 160X coverage by a paired-end strategy using Illumina HiSeq (paired-end read length: 2×75 nt) or NextSeq (paired-end read length: 2x∼75 nt) sequencers (Table S3). DNA-seq data were filtered on expected contaminants (ribosomal DNA, mitochondrial and bacterial genomes) then mapped on the *P.tetraurelia* MAC (ptetraurelia_mac_51.fa) or MAC+IES (ptetraurelia_mac_51_with_ies.fa) reference genome using Bowtie2 (v2.2.9 --local -X _500) 44,45._

### Bioinformatic analyses

Analyses were performed as previously described^8,39^. The IES annotation v1 (internal_eliminated_sequence_PGM_ParTIES.pt_51.gff3) was used for this study ^44^. These datasets are available from the ParameciumDB download section ^40^. The new version of ParTIES (v1.06 https://github.com/oarnaiz/ParTIES) was used for this study. The number of events was normalized by the number of mapped reads. R (v4.0.4) packages were used to manipulate annotations and to generate images (ggplot2 v3.3.5; GenomicRanges v1.42; rtracklayer v1.50).

## SUPPLEMENTAL FIGURES, FILES AND TABLES

**Supplementary figure S1. Survey of PolX in *P.aurelia* Species.**

(A) Analysis of the phylogeny of PolX identified in *Paramecium* species. PolXa (PTET.51.1.P0210235) sequence was blasted against the *Paramecium* genomes using pBlast (*Paramecium* DB^40^). Identified sequences were analysed together with Polλ using MUSCLE (EMBL-EBI web server). See Supplementary File S2 for multiple sequence alignment (MSA). The phylogeny highlights two main groups of PolX corresponding to the a-type (in blue) and c-type (in grey). All *P.aurelia* species, which were submitted to the two most recent WGDs, possess at least one representative of the a and c-type. (B) Left Panel, scheme of the nucleotide sequences of *POLX* open reading frames and the RNAi-inducing inserts cloned into the L4440 vector (Sequences of feeding vectors are presented in Supplementary File S1. Position 1 corresponds to the adenine of the starting codon ATG. IF1 sequences (in red) are specific of *POLX* paralogs, whereas IF3 (in blue) can target the paralogs of the last WGD. Right Panel, Percentage of survival of the sexual progeny using IF1 feeding inserts. (C) Survival of the sexual progeny after combinatory KD of *POLX* paralogs. The detail of 8 experiments is presented and was used to build the histogram presented in Figure 1C and Supplementary Figure 1B.

**Supplementary figure S2. Complementation assay using N-ter and C-ter flagged PolXa**

(A) Comparison of complementation efficiency using two differently Flag-tagged versions of PolXa. Western blot analysis of *FLAG-POLXa* and *POLXa-FLAG* expression levels in early autogamous cells (T5) subjected to *POLX* RNAi. The injection level, determined by qPCR, is indicated as copies per haploid genome (cphg). The percentage of survival of the sexual progeny is indicated for each transformant in the table below. (B) Replicates of the complementation experiments using FLAG-POLXa and FLAG-POLXc transformed cells. The injection level, determined by qPCR, is indicated (cphg). The percentage of survival of the sexual progeny is indicated for each transformant in the table below. The two transformants used for the immunolabelling experiment presented in the Figure 4A are surrounded.

**Supplementary figure S3. Detailed analysis of PGR genome-wide in control, PolXcd, PolXab and PolXabcd depleted cells.**

(A) MEND analysis of IES excision timing clusters. The number of reads was normalized by the number of IESs in each excision timing cluster. They are classified in five categories (see Figure 3A). (B) Histogram representation of IES retention score (IRS) distribution in PolXab and PolXcd depleted cells. (C) Screenshot of JBrowse (ParameciumDB) showing the position of the deletions (grey boxes) observed for three IES excision loci in PolX depleted cells. Reads are mapped on the MAC-51 genome. The IESs on the right and on the left have almost exclusively deletions on one side of the TA dinucleotide (red box at the top), whereas the IES in the middle shows reads carrying deletions on either side. (D) Genome-wide analysis of deletions around IES excision sites. The analysis was performed on the very early cluster of IESs (N=10994). The matrix represents the distribution of IESs (N=10994) according to the number of deletions that were counted at their −1 and +1 positions. (E) Logos of nucleotide frequencies found at IESs excision site carrying deletions. IESs were separated in six different groups, according to the presence (# reads > 5), the absence (# reads=0) of deletions, at the positions −1, +1 or both. The logos of base frequency for all very early IESs cluster is presented at the bottom as a control. (F) Analysis of the reads covering IES excision sites (using the MEND module of ParTIES) with or without directly compatible DNA ends. The figure represents relative proportion of reads and is restricted to very early excised IES, which were shown to be excised in the *POLX* KD.

**Supplementary figure S4. Western-blot control and detailed analysis of Flag signals relative to anlagen size in *POLXa*, *POLXc* and *POLXac* complemented cells.**

(A) Plot of Flag mean immunofluorescence intensity vs developing MAC size in *POLX* complemented cells. The size window, highlighted in grey, corresponds to the size of anlagen considered for quantification (see the boxplot analysis in figure 4A). The control of flagged proteins amount is presented in Supplementary Figure S2 for Flag-PolXa (672cphg) and Flag-PolXc (691cphg). (B) Plot of Flag mean immunofluorescence intensity *vs* developing MAC size in *POLXc* (998cphg) and *POLXac* complemented cells (clone1=698cphg and clone2=1328cphg). The size window (35-70μm^2^), highlighted in grey, correspond to the size of anlagen considered for quantification and was defined as previously^7^ (see the boxplot analysis in figure 4B). (C) Control of chimeric PolXa/c expression. The Flag-PolX signal, normalized by tubulin, and the percentage of sexual progeny survival is indicated below.

**Supplementary figure S5. Colocalisation of GFP-Dcl5 and endogenous Ku70a.**

*GFP-DCL5* transformed cells were analysed by immunolabelling of Ku70a using anti-Ku70 antibodies, at early time of autogamy (T5-T10), using a triton pre-extraction step before fixation.

**Supplementary file S1. Sequences of plasmids and oligonucleotides.**

**Supplementary file S2. Multiple sequence alignment of *P.Aurelia* PolX.**

**Supplementary Table S1. Sequencing Data**

**Supplementary Table S2. Quantification of Flag signal in immunolabelling experiment.**

## Notes

### Competing Interest Statement

The authors have declared no competing interest.

